# Perturbation of the human gut microbiome by a non-antibiotic drug contributes to the resolution of autoimmune disease

**DOI:** 10.1101/600155

**Authors:** Renuka R. Nayak, Margaret Alexander, Kye Stapleton-Grey, Carles Ubeda, Jose U. Scher, Peter J. Turnbaugh

## Abstract

The trillions of microorganisms (microbiota) found within the human gut play a critical role in shaping the immune system, yet these complex microbial communities are also highly sensitive to numerous environmental factors. While much of the focus to date has been on dietary intake, emerging data has begun to suggest that the use of pharmaceutical drugs, even those that are not considered to be antibiotics, can alter the human gut microbiota with unknown consequences for treatment outcomes. Here, we use a combination of *in vitro, in vivo*, and *ex vivo* methods to demonstrate that the first-line therapy for rheumatoid arthritis (RA), methotrexate (MTX), has off-target effects on the human gut microbiota, resulting in a significant growth advantage for drug-resistant Firmicutes over the Bacteroidetes, which tend to be more sensitive. Longitudinal analyses of the gut microbiotas of RA patients revealed that MTX-induced shifts in bacterial relative abundance are associated with improved drug response and transplant experiments in gnotobiotic mice show that these shifts lead to reduced inflammation. Together, these results suggest that the mechanism-of-action of non-antibiotic drugs may be due in part to off-target effects on the gut microbiota, while providing a critical first step towards explaining long-standing differences in drug response between patients.

## Introduction

The human gut microbiota consists of trillions of microorganisms that facilitate nutrient extraction from food^1^, avoidance of pathogen colonization^2^, and development of the host immune system^3^. Recent studies have highlighted how the gut microbiota is sensitive to multiple host and environmental factors^4,5^. While much of the focus has been on diet and host disease status, emerging data have demonstrated that pharmaceutical drugs can also impact the growth of human gut bacteria^6^. In a recent screen of 1,197 drugs, 24% were found to directly affect the *in vitro* growth of at least 1 of 40 human gut bacterial isolates^6^. However, the *in vivo* relevance and clinical significance of these off-target effects of non-antibiotic drugs remains to be determined.

Rheumatoid arthritis (RA) is an autoimmune disease of unknown etiology that affects between 0.5% to 1% of the population worldwide, leading to inflammation and destruction of joints as well as other organs^7^. Nearly all newly diagnosed RA patients are initiated on methotrexate (MTX)^8^. While about one third of patients demonstrate a favorable response to MTX^9^, needing little else for the management of their disease, the remaining two thirds require escalation of therapy with additional agents^10^. Furthermore, some patients may be intolerant of MTX, incurring side effects such as diarrhea, oral ulcers, liver toxicity or hair loss^11^. Thus, there is significant variability in patient response to MTX, with some patients experiencing great relief and others incurring little benefit from the drug^12^.

Although MTX is used in the treatment of RA and many other inflammatory diseases, it was originally developed as an anti-cancer therapy and was designed to target the “Achilles heel” of proliferating cells – the enzyme dihydrofolate reductase (DHFR)^13^. MTX is a structural analogue of folic acid, differing from this essential vitamin at only two sites. The DHFR enzyme converts folic acid into tetrahydrofolate (THF), which in turn serves as a carbon source for the production of purines and pyrimidines that are required for DNA synthesis and cell cycle progression^13^. Furthermore, THF provides carbon sources for pathways involving protein synthesis and lipid metabolism. For example, THF donates a methyl group to methionine, which then can be converted into S-adenosyl-methionine (SAM), a key molecule involved a myriad of pathways including protein synthesis, lipid methylation, and DNA methylation^14^. Thus, MTX targets a multitude of cellular functions downstream of DHFR, making it an effective anti-cancer drug.

Despite over 60 years of use in patients, the mechanism of action of MTX in modulating the immune system in RA and other rheumatologic disease is still not fully understood. MTX is used at far lower doses than those used in cancer^15^. Additionally, folic acid is frequently co-administered with MTX in order to reduce side effects in patients without an attendant loss in efficacy^14^, suggesting that MTX may act via other mechanisms besides DHFR inhibition. Here, we examine the impact of MTX on the gut microbiome, which in recent years has been shown to be a potent modulator of the host immune system^3,16,17^.

We reasoned that while MTX was originally developed to inhibit growth of human cells by targeting human DHFR, it may have “off-target” effects on human gut bacteria since DHFR is an enzyme that is conserved across all domains of life. Indeed, researchers have shown that MTX can bind DHFR homologues in *Escherichia coli* and *Lactobacillus casei*^18^. *E. coli* are resistant to any growth inhibitory effects of MTX due to the expression of a multi-drug efflux transporter *TolC*^19^. Overexpression of DHFR in a strain of *E. coli* that lacks *TolC* rescues it from the effects of MTX, suggesting that MTX interferes with DHFR in the cellular context^19^. Despite the discovery of this interaction 18 years ago, it remains unclear how generalizable this phenomenon is among other members of the human gut microbiota. A recent high-throughput screen^6^ confirmed these original findings and demonstrated that 12/40 strains were sensitive to MTX. We reasoned that this variation in the sensitivity of human gut bacteria to MTX may contribute to inter-individual variability in treatment outcomes.

Here, we use *in vitro* culturing, humanized mouse models, and the longitudinal analysis of human patients to demonstrate that the non-antibiotic drug MTX affects the structure and function of the human gut microbiome. Consistent with the prior literature in *E. coli*^6,19^, our data suggests that MTX acts via inhibition of bacterial DHFR, with downstream consequences for purine and pyrimidine biosynthesis. In treatment naïve RA patients, MTX induces a decrease in Bacteroidetes, consistent with our *in vivo* and *in vitro* data. However, this shift was only observed in drug responders, suggesting that MTX may act in part by modulating the gut microbiome. In support of this, MTX-altered microbiota from an RA patient (a drug responder) elicited less inflammation when transplanted into gnotobiotic mice. Together, these results demonstrate that drugs intended to target host pathways can have biologically relevant off-target effects on the gut microbiome, providing the first step towards addressing long-standing questions about the inter-individual variation in drug response and the integration of microbiome data for precision medicine.

## Results

### MTX has a dose-dependent impact on the gut microbiotas of humanized gnotobiotic mice

In order to determine whether MTX alters the community composition of the human microbiome *in vivo*, we utilized gnotobiotic mice colonized with a fecal aliquot from a healthy human male. We treated mice with 4 daily oral gavages of MTX at two doses, high (50 mg/kg) and low (1 mg/kg), selected to span the range of oral doses typically used in cancer^20^ and arthritis^21^, respectively. Vehicle controls were also included (n=3 mice/group). Daily stool samples were collected and processed for 16S rRNA gene sequencing to enable a “within-subjects” analysis of longitudinal changes in the gut microbiota for each mouse over the course of drug therapy. Endpoint samples were collected and analyzed from the small and large intestine (**Supplementary Table 1**).

MTX had a significant and dose-dependent impact on the gut microbiota. Total colonization based on 16S rRNA gene copy number (**Fig. 1a**) and microbial richness were comparable between treatment groups (**Fig. 1b**). In contrast, MTX treatment induced a significant shift in gut microbial community structure after a single day (ANOSIM, R=0.6049, *p*=0.004), and this effect persisted to day 4 of treatment (ANOSIM, R=0.7449, *p*=0.003) (**Fig. 1c**). There was no significant difference between groups prior to treatment (ANOSIM, *p*=0.109). Analysis with a second statistical method (PERMANOVA) confirmed these results (**Supplementary Table 2**). By day 4 of treatment, high-dose MTX significantly decreased the normalized abundance of the Bacteroidetes phylum (DESeq *p*_*adj*_=0.001, **Fig. 1d, Supplementary Fig. 1a**) compared to treatment with vehicle control. To take advantage of the “within-subjects” analysis of longitudinal changes, we used a generalized linear mixed-effects model to evaluate longitudinal changes^22^, and found that Firmicutes were significantly increased (slope=0.038, *p*=0.005) and Bacteroidetes were decreased (slope=-0.049, *p*=0.057). Eleven bacterial genera were differentially abundant comparing high-dose treatment to vehicle control (4 decreased and 7 increased, DESeq *p*_*adj*_ <0.01, day 4). At a finer level, we found that 82 ribosomal sequence variants (RSVs) were differentially abundant upon treatment with high-dose MTX (DESeq *p*_*adj*_<0.01, day 4; **Fig. 1e**, **Supplementary Table 3**). Low-dose MTX also significantly altered the gut microbiota, albeit to a lesser extent (20 were affected by low-dose MTX, DESeq *p*_*adj*_ <0.01, day 4, **Fig. 1e**, **Supplementary Table 4**). Fourteen RSVs were altered in both high- and low-dose MTX relative to vehicle-treated mice (**Fig. 1f**). These RSVs were altered in the same direction with either treatment, and the probability of this occurring by chance alone is exceedingly low (hypergeometric test, *p*=7.09 × 10^−5^). Together, these results demonstrate that MTX alters the human gut microbiome in a dose-dependent manner.

**Figure 1.**
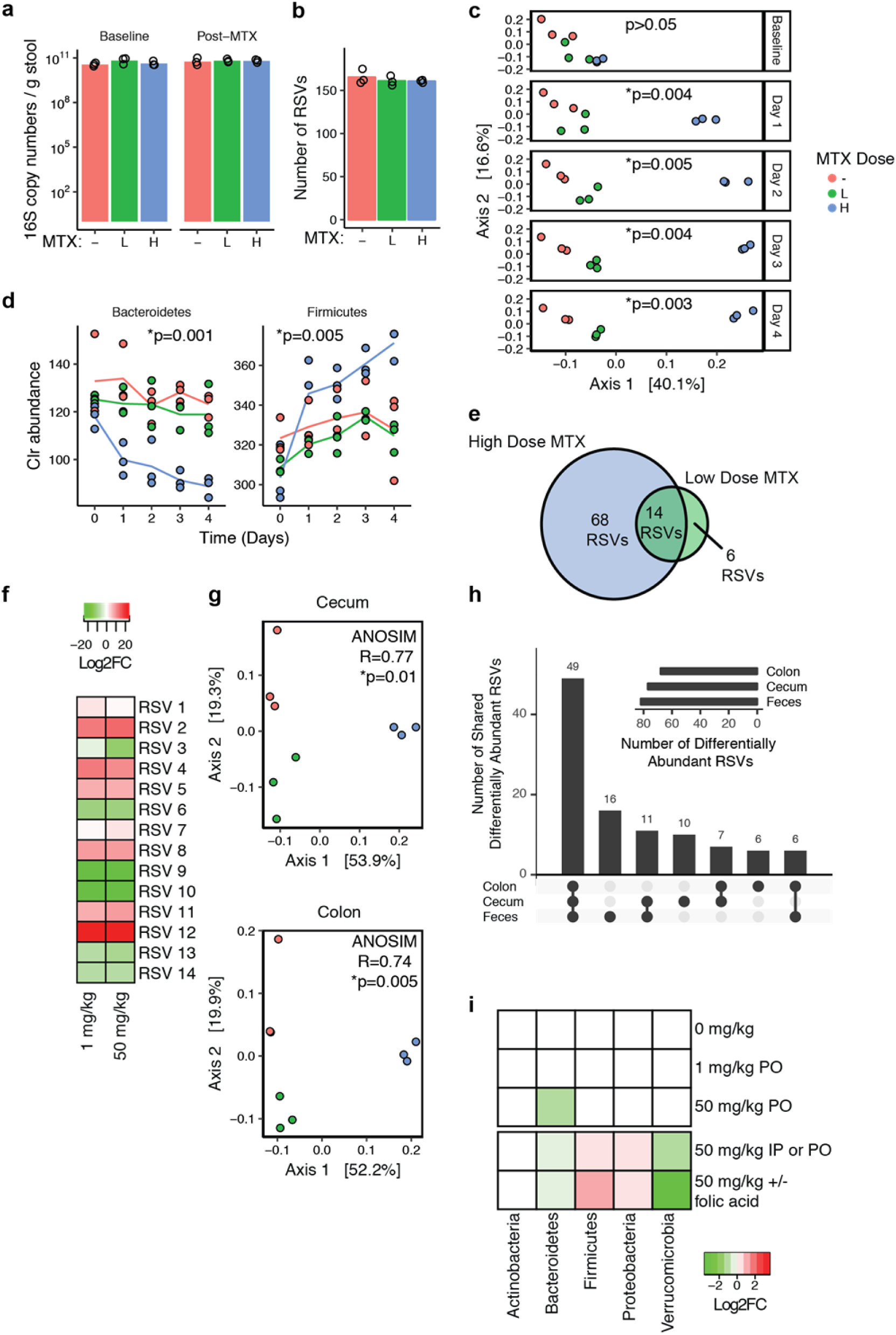
MTX alters the gut microbiomes of humanized mice. (**a**) Bacterial biomass as assessed by 16S copy number per gram of stool was quantitated using qPCR at baseline (Day 0) and after MTX treatment (Day 4). Treatments denoted by “-”, vehicle control; “L”, low-dose MTX (1 mg/kg); “H” high-dose MTX (50 mg/kg) (N=3 per treatment group). (**b**) Number of ribosomal sequence variants (RSVs) detected in treatment groups using 16S rRNA gene amplicon sequencing (N=3 per treatment group). (**c**) Community composition is altered with MTX treatment. Principal coordinates analysis (PCoA) of Bray-Curtis distances using clr transformed values at multiple time points after treatment. ANOSIM testing was performed comparing MTX at each time point. (**d**) The Bacteroidetes phylum significantly decreased with high-dose MTX treatment (DESeq *p*_*adj*_=0.001, comparing Day 4 vehicle vs. 50 mg/kg MTX). Firmicutes showed a time-dependent increase with treatment (p=0.005, generalized linear mixed modeling, see *Methods*). (**e**) Eighty-two and 20 RSVs were altered with 4 days of high- and low-dose MTX, respectively, relative to vehicle control. There were 14 RSVs that overlapped between these two sets. (**f**) Among the 14 RSVs that changed with both high- and low-dose treatment, all changed in the same direction. (**g**) Sites in the proximal GI tract also revealed changes in community composition with MTX treatment. (**h**) An UpSet plot of RSVs that were differentially abundant when comparing high-dose samples vs. vehicle controls in the cecum, colon and feces. The majority of RSVs that were differentially abundant in the feces were also differentially abundant in the cecum and colon. (**i**) A heatmap summarizing the significant phylum level changes observed in multiple gnotobiotic mouse experiments. MTX reproducibly induces a decrease in Bacteroidetes and Verrucomicrobia with a concomitant increase in Firmicutes and Proteobacteria (as assessed using DESeq comparing pre- and post-treatment time points).

To test if the changes observed in the fecal samples were reflective of changes in the small and large intestine, we sequenced samples from three sites (ileum, cecum, and colon). We were unable to detect significant changes in the ileum. We suspect this may be due to limited recovery because the ileum has a lower bacterial burden than the distal GI tract^23^, and we subsequently obtained less sequencing coverage (**Supplementary Table 1**). In contrast, the cecum and colon demonstrated significant changes in community composition (cecum: ANOSIM R=0.7695, *p*=0.01; colon: ANOSIM R=0.7366, *p*=0.005) (**Fig. 1g**). Twenty-one and 23 RSVs were significantly different in the cecum and colon with low-dose treatment, respectively, and 77 and 68 RSVs were significantly different in the cecum and colon with high-dose treatment, respectively (DESeq *p*_*adj*_ <0.01, **Supplementary Table 5, 6**). The majority of the RSVs (60% or 49/82) that were significant in the fecal samples with high-dose treatment were also significant in the cecal and colon samples (**Fig. 1h**), consistent with our prior data that fecal compositional trends reflect changes within the gastrointestinal (GI) tract^24^.

MTX is administered subcutaneously or intravenously in some patients and undergoes enterohepatic circulation, returning a portion of the systemic drug to the gastrointestinal tract^25,26^. To test the reproducibility of our findings given different dosing methods and in the context of a disease-associated microbiome, we compared oral (PO) administration of MTX to intraperitoneal (IP) injection in humanized gnotobiotic mice colonized with stool from a MTX-naïve RA patient (n=6 adult female C57BL/6J recipients; 50 mg/kg daily for 2 days). Analysis of gut microbial community structure based on 16S rRNA gene sequencing revealed that there was no significant difference in the gut microbiotas of mice administered PO *vs.* IP MTX (ANOSIM R=0.111, *p*=0.4). When pooling both treatments and comparing pre- vs. post-treatment samples, both groups experienced a significant effect on community composition (ANOSIM R=0.4756, *p*=0.019; PERMANOVA *p*=0.001; comparing Day 0 vs. Day 2). The Bacteroidetes phylum decreased in abundance in both PO and IP treated mice, replicating the effects we saw in our initial experiment (-1.45 fold-change, *p*_*adj*_<10^−5^, DESeq) (**Fig. 1i, Supplementary Fig. 1b**). We detected a significant increase in two other phyla, Firmicutes (1.65 fold-increase; *p*_*adj*_=0.007) and Proteobacteria (1.35 fold-increase; *p*_*adj*_ =0.01), accompanied by a significant decrease in Verrucomicrobia (2.54 fold-decrease; *p*_*adj*_ =7.27 × 10^−4^) (**Fig. 1i**). We also detected a significant impact of MTX at finer taxonomic levels: 10 genera were significantly changed (4 increased and 6 decreased, *p*_*adj*_ <0.05, DESeq) and 51 RSVs were significantly altered (24 increased and 27 decreased, *p*_*adj*_ <0.05, DESeq, **Supplementary Table 7**). Together, these results confirm that MTX alters the gut microbiome regardless of the route of delivery.

We next asked whether folic acid rescues the effects of MTX on the microbiome, since this drug is used to rescue the effects of MTX toxicity in humans^27^. We treated mice either with oral MTX 50 mg/kg or with MTX and folic acid in approximately a 1:1 molar ratio at 50 mg/kg daily for 2 days. Community composition was altered in both treatment groups to similar extents and we did not find a significant difference between the two groups (ANOSIM R=0.7401, *p*=0.1), suggesting that folic acid at the dose given in our study was not sufficient to rescue the effects of MTX on the microbiome. When comparing pre- and post-treatment samples using DESeq, we saw reproducible trends at the phylum level: a decrease in Bacteroidetes and Verrucomicrobia (fold changes of-1.44 and -12.47 with *p*_*adj*_ =3.69 × 10^−6^ and *p*_*adj*_ =3.7 × 10^−21^, respectively), and an increase in Firmicutes (1.76 FC; *p*_*adj*_ =4.56 × 10^−8^) and Proteobacteria (1.39 FC; *p*_*adj*_ =0.001) (**Fig. 1i, Supplementary Fig. 1c**). Twenty genera (10 increased and 10 decreased, *p*_*adj*_ <0.05, DESeq) and 67 RSVs (32 increased and 35 decreased, *p*_*adj*_ <0.05, DESeq, **Supplementary Table 8**) were affected by MTX.

Taken together, these experiments demonstrate that MTX reproducibly alters the human gut microbiota in gnotobiotic mice, with reproducible effects across multiple delivery routes and co-therapies. Next, we sought to test if MTX directly impacts the growth of human gut bacteria or reflects a downstream consequence of this drug on host tissues.

### Methotrexate directly inhibits the growth of human gut bacteria

High-dose MTX acts by inhibiting dihydrofolate reductase (DHFR)^13^, an enzyme conserved across all 3 domains of life^28^. Given the importance of DHFR for bacterial growth across multiple phyla^29^, we hypothesized that MTX could act as a broad-spectrum antibiotic. To test for a direct impact of MTX on bacterial cell growth, we incubated MTX with a panel of 43 bacterial isolates from 6 phyla, 40 of which are commonly found in the human gut microbiota (**Supplementary Table 9)**. The exceptions include *Bacteroides acidifaciens* (from mice), and *Delftia acidovorans* and *Bacillus subtilis* 168 (from soil). The combined average relative abundance of these bacterial isolates is 43%^30^ of the human gut microbiota (**Supplementary Fig. 2**; *Methods*). Each isolate was incubated with a gradient of MTX ranging from 0-900 µg/ml (0-2 mM) and bacterial growth in rich media was quantified by optical density. The concentration required to prevent ≥90% growth (MIC_90_) ranged three orders of magnitude: from 2 μM to 2 mM (**Fig. 2a**). While 11 of 45 isolates (24%) were resistant to the maximum concentration tested, 34 isolates (76%) were variably sensitive to the growth inhibition induced by MTX. On average, Bacteroidetes tended to be sensitive to the antimicrobial effects of MTX relative to the other phyla (**Fig. 2b**, *p*=0.005, Wilcoxon rank-sum test).

**Figure 2.**
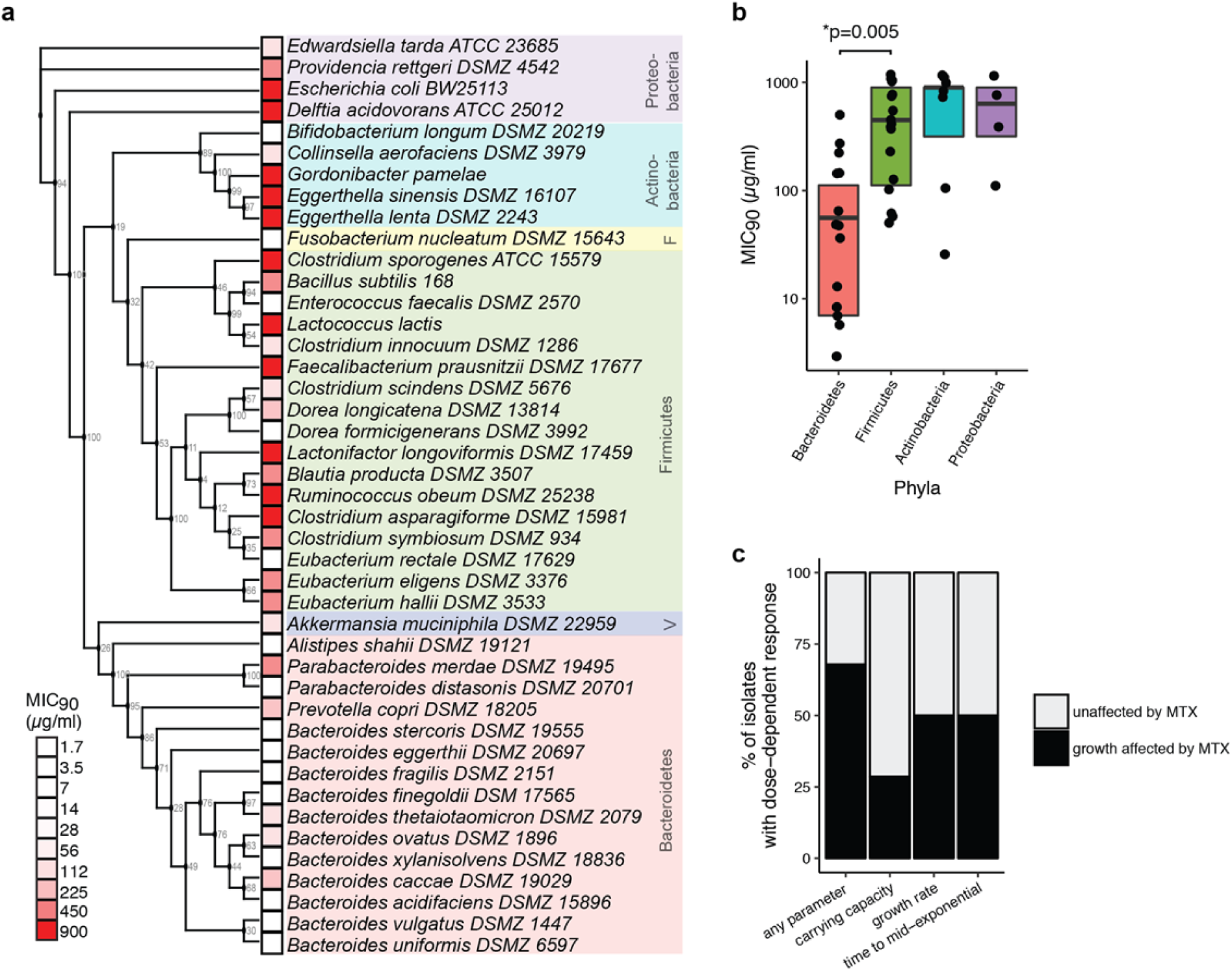
MTX directly affects growth of human gut bacteria. (**a**) A diverse panel of 43 isolates were incubated with varying concentrations of MTX and the minimal inhibitory concentration (MIC_90_) was measured. F, Fusobacteria; V, Verrucomicrobia. A maximum likelihood phylogenetic tree using 16S rRNA gene sequences for each organism was constructed, with bootstrap values from 100 iterations shown at each branch point. (**b**) The MICs of various isolates spanning 4 major phyla. Bacteroidetes tended to be sensitive relative to other phyla when tested *in vitro* (*p*=0.005, Wilcoxon rank-sum test). The top and bottom hinges in the boxplot correspond to the first and third quartile, respectively, and the horizontal lines denotes the median. (**c**) Carrying capacity, growth rate and lag phase parameters were affected in a dose-dependent manner by MTX among a significant proportion of bacteria.

Nineteen (42%) of the MIC_90_ values were at or below the predicted concentration of MTX in the proximal GI tract (250 μM, 113 μg/ml, *Methods*)^31^, suggesting that MTX is capable of inhibiting growth of these isolates at physiological concentrations. Among the remaining 26 isolates (58%) whose MIC_90_ are above the predicted concentration, we asked whether MTX could still significantly alter the growth of these isolates by examining carrying capacity, growth rate, and the duration of lag phase (*i.e.* time to mid-exponential growth). We found that 19 of the remaining 26 (73%) isolates showed changes in at least one of these three growth parameters in a dose-dependent manner (**Fig. 2c, Supplementary Fig. 3**, see *Methods*). Taken together, our results show that 38 of the 45 (84%) tested isolates exhibit either growth inhibition or alterations in growth curve parameters upon exposure to MTX at concentrations that are predicted to be found in the human gut. These results raised the question of whether or not MTX acts through the canonical mechanism-of-action established in eukaryotic cells or if it has off-target effects on other aspects of bacterial physiology.

### Methotrexate impacts gut bacterial purine and pyrimidine biosynthetic pathways

Prior studies have revealed that MTX can directly inhibit bacterial DHFR^18^; however, the downstream consequences of this inhibition for bacterial physiology remain unexplored. To globally profile the impact of bacterial DHFR inhibition, we used transcriptomics (RNA-Seq) to search for differentially expressed transcripts in the presence or absence of MTX. Based on the extensive literature in human cells^32,33^, we anticipated that the inhibition of DHFR by MTX would lead to changes in the expression of purine and pyrimidine pathways, which rely on folate as a co-factor for multiple key reactions. We selected isolates with varying sensitivity to the growth-inhibitory effects of MTX: *Clostridium asparagiforme* DSMZ 15981 (MIC_90_ > 900 μg/ml), *Clostridium sporogenes* ATCC 15579 (MIC_90_ > 900 μg/ml), *Clostridium symbiosum* DSM 934 (MIC_90_ = 450 μg/ml), and *Bacteroidetes thetaiotaomicron* DSMZ 2079 (MIC_90_ = 112 μg/ml). Bacteria were treated with MTX 100 μg/ml or vehicle control (DMSO) for 30 minutes during mid-exponential phase (n=3 per treatment group per isolate).

MTX induced a profound shift in gene expression in the isolate that was most sensitive to MTX, *B. thetaiotaomicron*, affecting 83% of genes in the transcriptome (**Fig. 3a**, 3,735 genes at FDR<0.2, DESeq). In contrast, two less sensitive isolates, *C. sporogenes* and *C. symbiosum*, exhibited a more defined shift in gene expression in response to MTX (**Fig. 3a**, 21 and 55 genes, FDR<0.2, DESeq, respectively). Although *C. asparagiforme* did not show significant growth inhibition with MTX (**Fig. 2a**), it had a robust transcriptional response to the drug, with 468 genes (8% of expressed transcripts) exhibiting differential expression at 30 minutes. These results demonstrate that gut bacterial strains vary in their response and sensitivity to MTX at the transcriptional level. Surprisingly, some drug resistant bacteria still show a significant transcriptional response to the drug, consistent with our prior observations that MTX can impact bacterial physiology even below the MIC_90_.

**Figure 3.**
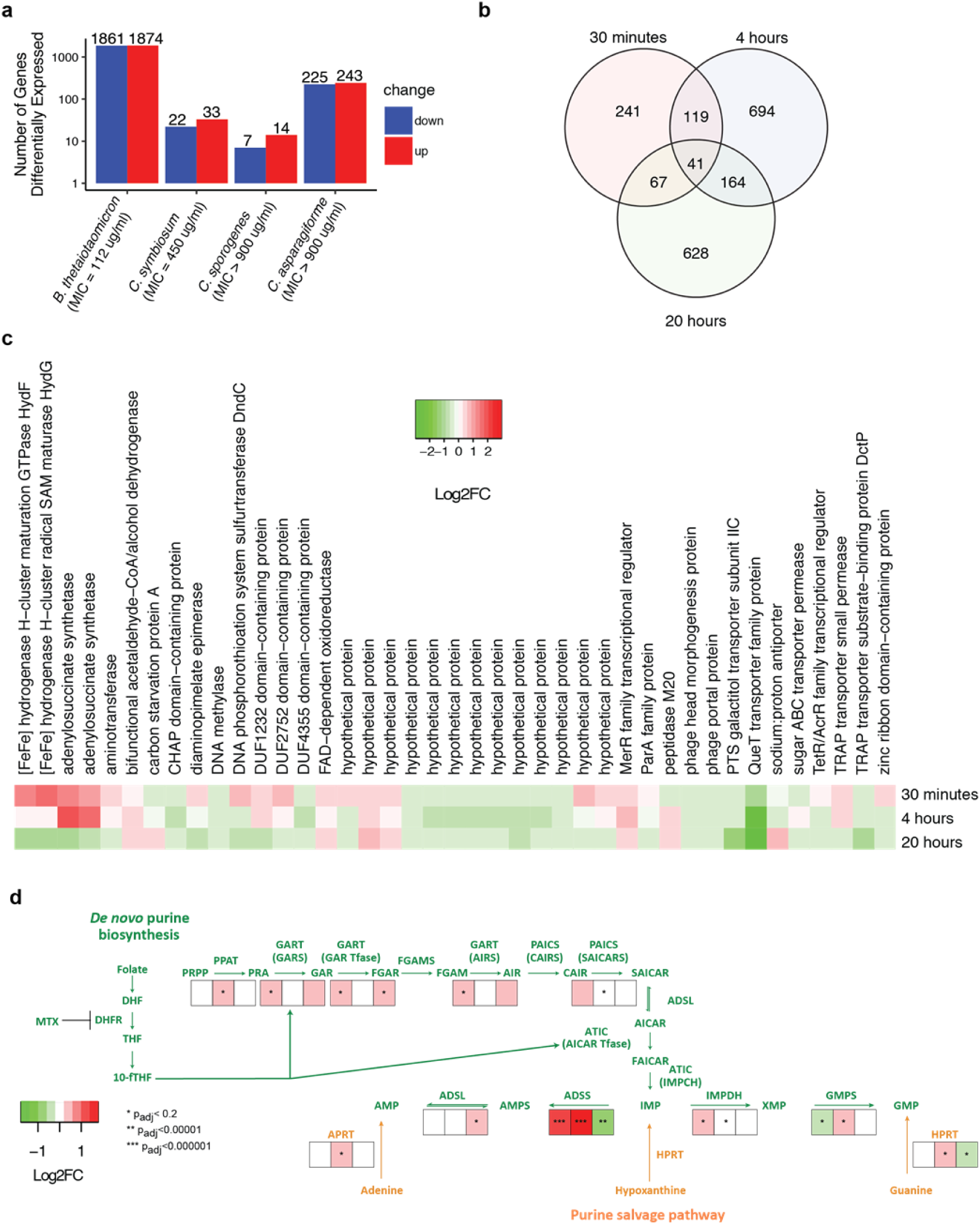
MTX affects transcriptional pathways associated with purine and pyrimidine metabolism in human gut bacteria. (**a**) A variable number of transcripts, as determined by RNA-Seq, were differentially expressed (FDR<0.2, DESeq) upon 30 minutes of MTX 100 μg/ml (compared to vehicle control) in 4 bacterial isolates with varying sensitivity to the growth inhibitory effects of the drug (n=3 per treatment). *B. thetaiotomicron*, which is sensitive to the growth inhibitory effects of MTX, exhibited a profound alteration in transcriptional response. *C. asparagiforme*, which is resistant to the growth inhibitory effects of MTX, also manifested a transcriptional response. (**b**) We performed a time course analysis on *C. asparagiforme*, which was profiled at 30 minutes, 4 hours and 20 hours of treatment (n=3 per treatment group per time point). The Venn diagram shows the number of transcripts that were differentially expressed with each treatment duration. (**c**) A heatmap of the 41 transcripts that were differentially expressed (FDR<0.2) at all three time points in *C. asparagiforme*. (**d**) Multiple enzymes involved in purine metabolism^36^ were affected at some point during the time course study that was performed on *C. asparagiforme* (heatmap of log2 fold change values for enzymes shown, with asterisks indicating *p*_*adj*_ values). See Supplementary Table 13 for a description of substrate and enzyme symbols.

To gain insight into the metabolic pathways most impacted by MTX, we used the KEGG database to search for differentially expressed metabolic pathways and modules. Both purine and pyrimidine metabolism were significantly changed in *C. asparagiforme* and *B. thetaiotaomicron*. While *B. thetaiotaomicron* exhibited multiple pathway enrichments (57 pathways enriched among transcripts with FDR<0.2), purine and pyrimidine metabolism were among the top 10 pathways (9^th^ and 5^th^, respectively) when ranked by *p*-value (*p*<0.05 with BH adjustment), and these enrichments were insensitive to the FDR threshold used in our analyses (**Supplementary Table 10**). *C. asparagiforme* exhibited enrichment of 23 pathways, and purine and pyrimidine metabolism were 2^nd^ and 11^th^ among this list (**Supplementary Table 11**). These results suggest that MTX can exert wide-ranging effects on numerous pathways in sensitive bacteria like *B. thetaiotaomicron*, or have more specific effects on metabolic pathways in bacteria like *C. asparagiforme*.

To assess the dynamics of sub-MIC_90_ MTX response, we performed a time course experiment on *Clostridium asparagiforme* (MIC_90_ >900 μg/ml), comparing transcriptional profiles at 30 minutes, 4 hours and 20 hours post drug exposure to vehicle controls. We found that 1,018 genes were affected at 4 hours and 900 affected at 20 hours (**Fig. 3b**; FDR<0.2). Consistent with our original analyses, purine and pyrimidine metabolism continued to be among the pathways that were affected at 4 and 20 hours of treatment in *C. asparagiforme* (**Supplementary Table 11**). Of the 41 transcripts that were differentially expressed at all three timepoints (FDR<0.2, DESeq), 21 consistently changed in the same direction (5 upregulated, 16 down-regulated), whereas 20 demonstrated more complicated dynamics (**Fig. 3c**).

Because MTX is thought to target multiple enzymes, including DHFR^34^ and AICAR transformylase^35^ which are both involved in purine metabolism, we examined the homologs of enzymes involved in purine metabolism in *C. asparagiforme*. While we did not find that DHFR or AICAR transformylase were differentially expressed, we found that multiple enzymes either upstream or downstream of these genes were differentially expressed. Enzymes involved in *de novo* purine biosynthesis as well as the salvage pathway for purine synthesis showed differential expression (**Fig. 3d**). For example, adenylosuccinate synthase (ADSS), which is involved in converting inosine monophosphate (IMP) into adenylosuccinate (AMPS)^36^, is significantly upregulated. These findings further support the hypothesis that MTX targets bacterial enzymes involved in purine metabolism; however, it remains possible that other pathways would be more dramatically affected in later stages of drug exposure.

Taken together, these results suggest that while MTX was developed as a drug to target human cells, it has off-target effects on pathways for purine and pyrimidine metabolism in bacterial cells. MTX inhibits the growth of several gut bacterial isolates in a dose-dependent manner. Transcriptionally, it targets pathways that are reliant on folate, including purine and pyrimidine metabolism, but can also affect more widespread pathways as seen in *B. thetaiotaomicron* and the 20-hour time point in *C. asparagiforme*. Our data from *C. asparagiforme* demonstrates that even when MTX does not affect the growth of the bacteria, it can still affect transcriptional pathways that depend on folic acid.

### Clinical relevance of the interaction between MTX and the human gut microbiome

A major caveat of our isolate-based studies is that human gut bacteria may not be directly sensitive to MTX in the context of a complex gut microbial community or in the physiochemical conditions found within the human gastrointestinal tract. To address this concern, we turned to *ex vivo* incubations of human distal gut microbiomes and studies of patient cohorts. First, we examined whether MTX affects growth in *ex vivo* stool samples obtained from 30 MTX-naïve patients. We treated fecal suspensions with MTX 100 μg/ml or vehicle control and measured growth by optical density over 48 hours. We found that MTX inhibits growth of these complex *ex vivo* communities (**Fig. 4a**). Specifically, we found that MTX decreases the carrying capacity (*p*<10^−5^, paired Student’s *t*-test, **Fig. 4b**) and increases the time to mid-exponential growth (*p*<0.001, paired Student’s *t*-test, **Fig. 4c**) in *ex vivo* samples, supporting the hypothesis that MTX can have growth inhibitory effects on patient microbiomes *ex vivo*.

**Figure 4.**
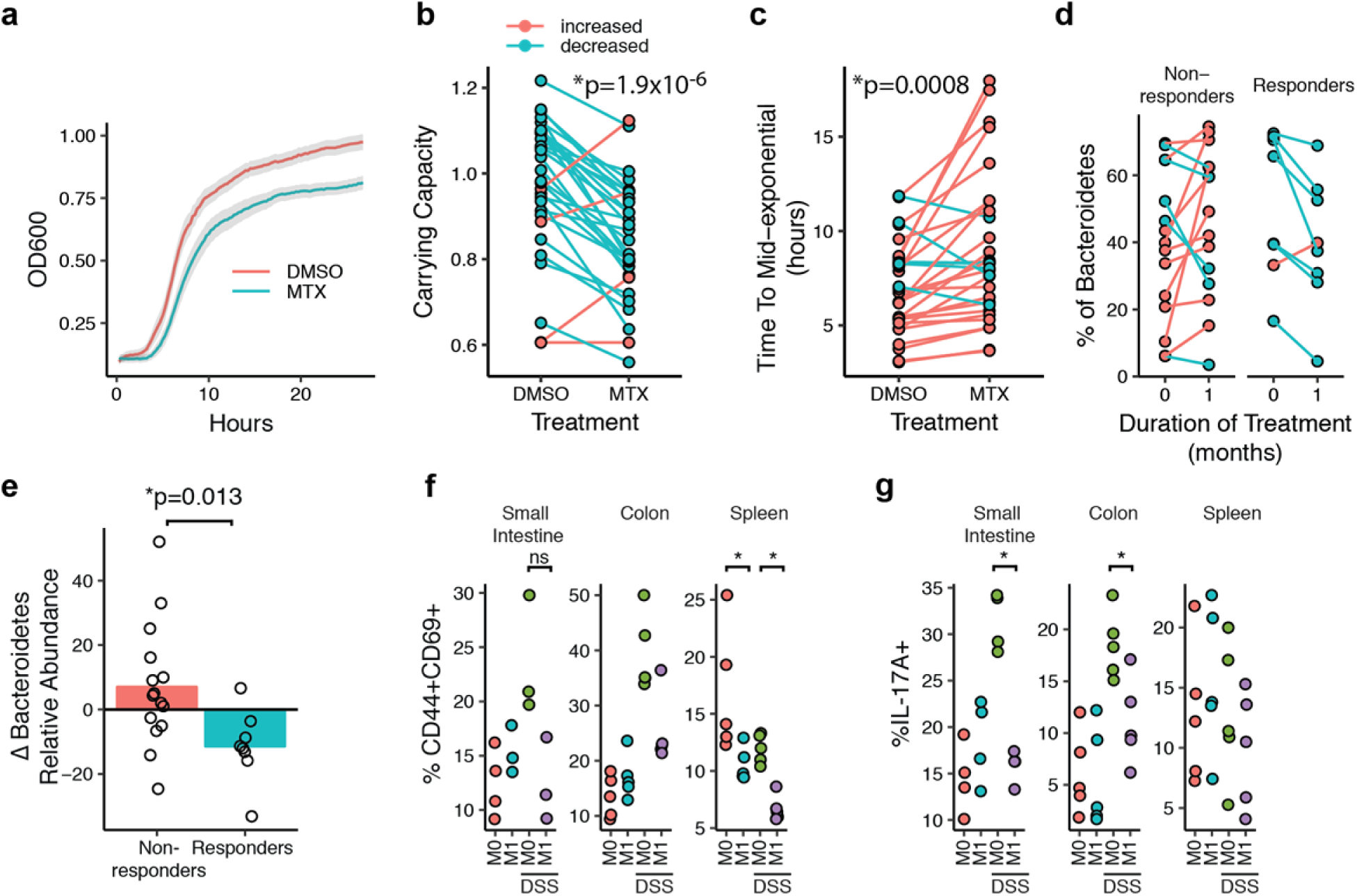
Human gut bacteria from rheumatoid arthritis (RA) patients are affected by MTX *ex vivo* and *in vivo*, and MTX alters their inflammatory potential. (**a**) Growth of fecal suspensions from treatment-naive RA patients (n=30) was measured using optical density in the presence of MTX 100 μg/ml or DMSO. The average growth curves among the 30 patients are shown for each treatment. Shaded areas represent ±SEM. (**b**) Carrying capacity was significantly decreased among the 30 patient fecal suspensions *ex vivo* (paired Student’s *t*-test). (**c**) Time to mid-exponential was significantly increased among the 30 patient fecal suspensions *ex vivo* (paired Student’s *t*-test). (**d**) Fecal samples from 23 RA patients was subjected to 16S sequencing before treatment with MTX and 1 month after treatment. Patients that demonstrated a favorable clinical response to MTX showed a decrease in relative abundance of Bacteroidetes compared to non-responders. (**e**) There is a significant decrease in the relative abundance of Bacteroidetes with MTX treatment among responders compared to non-responders (*p*=0.013, Wilcoxon rank-sum). (**f**) Percentage of CD3+CD4+ T cells with CD44+CD69+ staining (activated T cells) from the small intestine and colon lamina propria and spleen of mice transplanted with microbiota from a patient before MTX treatment (“M0”) and 1 month after treatment (“M1”) (N=3-5 per treatment group). Mice were also treated with dextran sodium sulfate (“DSS”) in their drinking water. (**g**) Percentage of IL-17A+ cells in the CD3+CD4+ T cell compartment as determined by flow cytometry in the small intestine and colon lamina propria and spleen (N=3-5 per treatment group). Panels f,g: \**p*<0.05, 2-tailed Student’s *t*-test.

We next asked whether these findings extended to the *in vivo* setting in RA patients. We performed 16S rRNA gene sequencing on stool samples collected at baseline and 1 month after treatment initiation from 23 MTX-naïve patients. Consistent with prior reports^24,37-39^, we detected a strong signal of individuality in gut microbial community structure (R=0.9152, *p*=0.001, ANOSIM). Although there were marked shifts in community structure within each subject following treatment (**Supplementary Fig. 4**), we were unable to detect a reproducible shift across the entire patient cohort in response to MTX (ANOSIM, R=-0.09, *p*=0.888), consistent with prior studies^40^. We anticipated an individualized microbial community response to MTX given inter-individual differences in the gut microbiota prior to therapy coupled to the extensive strain-level variation in drug sensitivity identified in this study.

These results led us to hypothesize that differences in the gut microbial response to MTX could explain some of the differences in clinical response. Given our relatively small sample size, we used a binary classification of drug response defined as a decrease in DAS28^41^ of at least 1.8 and continued use of oral MTX without addition of other disease-modifying anti-rheumatic drugs (DMARDs) or biologics. Using these criteria, our cohort was split between 15 MTX non-responders (MTX-NR) and 8 responders (MTX-R). The baseline disease activity parameters between these two groups did not differ (**Supplementary Table 12**). Among the 8 MTX-R, we observed a significantly decreased relative abundance of Bacteroidetes compared to MTX-NR (*p*=0.01, Wilcoxon rank-sum test, **Fig. 4d, 4e**). This decrease in Bacteroidetes recapitulated results we observed in our *in vitro* and humanized gnotobiotic mouse studies and led us to ask whether drug-induced shifts in the human gut microbiota contribute to the resolution of autoimmune disease.

### MTX-altered microbiota dampen host immune responses

To determine whether the anti-inflammatory effects of MTX are mediated in part via the microbiome, we examined the functional impact of MTX-altered microbiota on mucosal and peripheral T cell populations in gnotobiotic mice. We identified the MTX-responder among our patient cohort with the greatest decrease in Bacteroidetes and transferred fecal microbiota obtained before (M0) and 1 month after MTX treatment (M1) from this donor into C57BL/6J germ-free mice (N=10 per donor sample, 20 female mice total, **Supplementary Fig. 4b**). Half of the animals in each group were challenged with an inflammatory trigger using 2% dextran sodium sulfate (DSS) (N=5 per donor sample), a chemical added to the drinking water used to induce rapid and robust colitis. Since RA pathophysiology is associated with a dysregulated T cell response^42^, we focused on the T cell compartment in the spleen and the intestinal lamina propria of the small intestine and colon using flow cytometry. In unchallenged mice, we found that splenic populations of activated (CD44+CD69+) T cells were decreased in mice transplanted with MTX-altered microbiota (M1) compared to M0 microbiota (10.6% vs. 16.8%; *p*=0.04, Student’s *t*-test; **Fig. 4f**). We did not observe significant differences in mucosal T cell populations (Th17, Th1 or T regulatory cells) or in myeloid cells in unchallenged mice (**Supplementary Fig. 4c**).

In mice challenged with DSS, similar extents of weight loss, clinical colitis, and colonic shortening were observed between transplant groups (**Supplementary Fig. 4d**). Examination of immune cell populations confirmed that DSS induced an inflammatory response (**Fig. 4f,g**). T cell activation in the spleen was decreased in mice colonized with M1 compared to M0 microbiota (6.7% vs. 12%; *p*=0.0001, Student’s *t*-test; **Fig. 4f**), similar to what was observed in the unchallenged state. Furthermore, splenic T cells expressing IFN-γ were reduced (3.5% vs. 4.8%; *p*=0.01, Student’s *t*-test; **Supplementary Fig. 4c**). In mice harboring M1 microbiota, Th17 T cells were relatively decreased in the small intestine (15.8% vs. 31.4%; *p*=0.0001, Student’s *t*-test) and colon (11.1% vs. 18.5%; *p*=0.01, Student’s *t*-test) (**Fig. 4g**), suggesting that MTX treatment leads to a reduction in the Th17-inflammatory potential of human gut microbiota. Overall, these findings suggest that MTX-induced shifts to the microbiota reduce its inflammatory potential and may contribute to resolution of autoimmune disease.

## Discussion

While MTX was developed to target human dihydrofolate reductase (DHFR), it was not designed to be specific for the human enzyme, leading us to ask whether this non-antibiotic drug may have direct, off-target effects on bacteria. Previous studies demonstrated that community composition is altered in rheumatoid arthritis patients relative to healthy individuals^43^ and that this alteration is “reversed” with treatment^44^. But in these patient studies, it is unclear whether the changes to the microbiome are a result of drug treatment, concomitant treatment with other medications, improvement of the disease process, reduction of inflammation, or other confounding factors that may be present in observational cohort studies. Our studies in disease-free gnotobiotic mice, coupled to *in vitro* and *ex vivo* results, demonstrate that MTX directly exerts growth inhibitory effects on a diverse panel of human gut bacterial isolates with downstream consequences for bacterial physiology and the expression of genes involved in purine and pyrimidine metabolism. Thus, our findings extend our knowledge of the effects of MTX on the host to include its effects on the gut microbiome.

These results provide a critical foundation to ask whether MTX may exert part of its anti-inflammatory effects via the gut microbiome. Although MTX has been used for several decades to reduce inflammation in patients with autoimmune disease, the mechanisms by which it does so are not fully understood. A leading hypothesis is that MTX directly inhibits an enzyme called AICAR transformylase, resulting in an increase in extracellular levels of adenosine, a potent anti-inflammatory mediator^14,45^. Other possible mechanisms that have been investigated suggest that MTX may affect cytokine production by immune cells, such as by decreasing the production of the proinflammatory cytokine TNFα^46^ or increasing the production of the α-inflammatory cytokine IL-10^47^. But it remains unknown whether MTX reduces inflammation by altering the gut microbiome, which has been shown to be a potent modulator of the host immune system. Importantly, there is a precedent for a drug’s therapeutic effects being mediated via the microbiome in the context of the treatment of diabetes with metformin^48,49^.

We provide evidence for such a mechanism by transferring microbiota from a MTX-treated RA patient donor (obtained before and 1 month after treatment) into germ-free mice. We found that MTX results in a decrease in Bacteroidetes, permitting members of other bacterial phyla to expand in relative abundance, and that these shifts alter the inflammatory potential of the microbiome. In particular, we found a reduction in Th17 cells in the mucosa and a reduction in activated T cells and IFN-γ+ T cells in the periphery. These cell types are thought to play key roles in RA pathogenesis^42^, and prior studies suggest that MTX decreases IL-17 levels in RA patients^50^. By uncoupling the effect of MTX on the host from its effect on the gut microbiome, these gnotobiotic results provide causal evidence suggesting that MTX exerts its anti-inflammatory effects in part by reducing the ability of the gut microbiome to contribute to an inflammatory response. Our findings advance the current understanding of the pharmacodynamics of MTX and provide a critical foundation to further investigate the microbial effectors of these immune responses.

The impact of MTX on bacterial biomass are not as pronounced as seen with many broad-spectrum antibiotics^51^; however, our data shows that MTX can have effects on bacterial physiology at a sub-inhibitory concentration. These results indicate that we are likely underestimating the effect of MTX (and possibly other drugs^52^) on the gut microbiome when examining community composition by DNA sequencing alone. As expected, bacteria sensitive to the growth inhibitory effects of MTX, such as *B. thetaiotaomicron*, exhibited profound transcriptomic changes. Surprisingly, we found that *C. asparagiforme*, which is resistant to the growth inhibitory effects of MTX, experienced pronounced changes at the transcriptional level. Thus, *in vivo*, MTX may not only affect community composition, but it may also have broader effects on transcriptional and metabolic programs of human gut microbiota. Further studies are needed to examine transcriptional and metabolic changes to the human gut microbiome induced by MTX in both humans and mouse models.

While we did not observe a consistent effect of MTX on community composition among all RA patients when examining 16S rRNA gene sequencing-based community composition, this may be because there is heterogeneity among RA patients with respect to the drug disposition^53^, genetic risk factors^54^, and/or microbiome composition^44^. Our data demonstrates that gut bacterial strains and intact microbial communities can vary dramatically in their drug sensitivity. The underlying determinants of this variability remain to be investigated and are likely multifactorial. Of note, responders tend to experience a greater decrease in Bacteroidetes upon initiation of MTX relative to non-responders. Additionally, the transcriptional response to MTX *in vivo* can be greater than detected by 16S rRNA gene sequencing alone.

Our data suggests that the interaction between MTX and the gut microbiome is broadly relevant to patients even given differences in dosing route and adjuvant therapies. Both intraperitoneal dosing and oral dosing impacted the gut microbiome, likely due to extensive enterohepatic circulation^25,26^. Similar microbiome effects were observed with folic acid supplementation, which is commonly given in RA patients, suggesting that the dose administered does not markedly rescue *in vivo* bacterial growth inhibition. Follow-on longitudinal analysis in patients given a variety of treatment regimens would help to further clarify whether or not the impact of MTX on the gut microbiome depends at all on common clinical confounders.

In conclusion, our results emphasize the importance of taking a broader view of toxicology that encompasses the unintended consequences of non-antibiotic drugs for our associated microbial communities. Our studies demonstrate the utility of integrated studies *in vitro*, in gnotobiotic mice, *ex vivo*, and in drug naïve patients to begin to elucidate the causality and mechanism for these complex drug-microbiome-host interactions. Remarkably, the observed drug-induced changes in microbial community structure were associated with patient response, providing a useful biomarker for accelerating the stable initiation of therapy and a first step towards determining which bacterial taxa contribute to or interfere with treatment outcomes.

## Acknowledgments

Special thanks to Lindsey Criswell, Kathy Lam, Mary Nakamura, and Katie Pollard for comments on the manuscript. We are indebted to Jessie Turnbaugh and the Gnotobiotic Core for technical assistance as well as the Institute for Human Genetics for help with RNA sequencing. We thank Jordan E. Bisanz and Elizabeth N. Bess for their assistance with sequencing methods.

## Author contributions

Conceptualization, R.N. and P.J.T.; Investigation, R.N., M.A., K.S.G; Data Analysis, R.N.; Clinical Samples Acquisition and Sequencing, J.S. and C.U.; Writing – R.N. and P.J.T.; Funding Acquisition, R.N. and P.J.T.

## Supplementary Figures & Legends

**Supplementary Figure 1.**
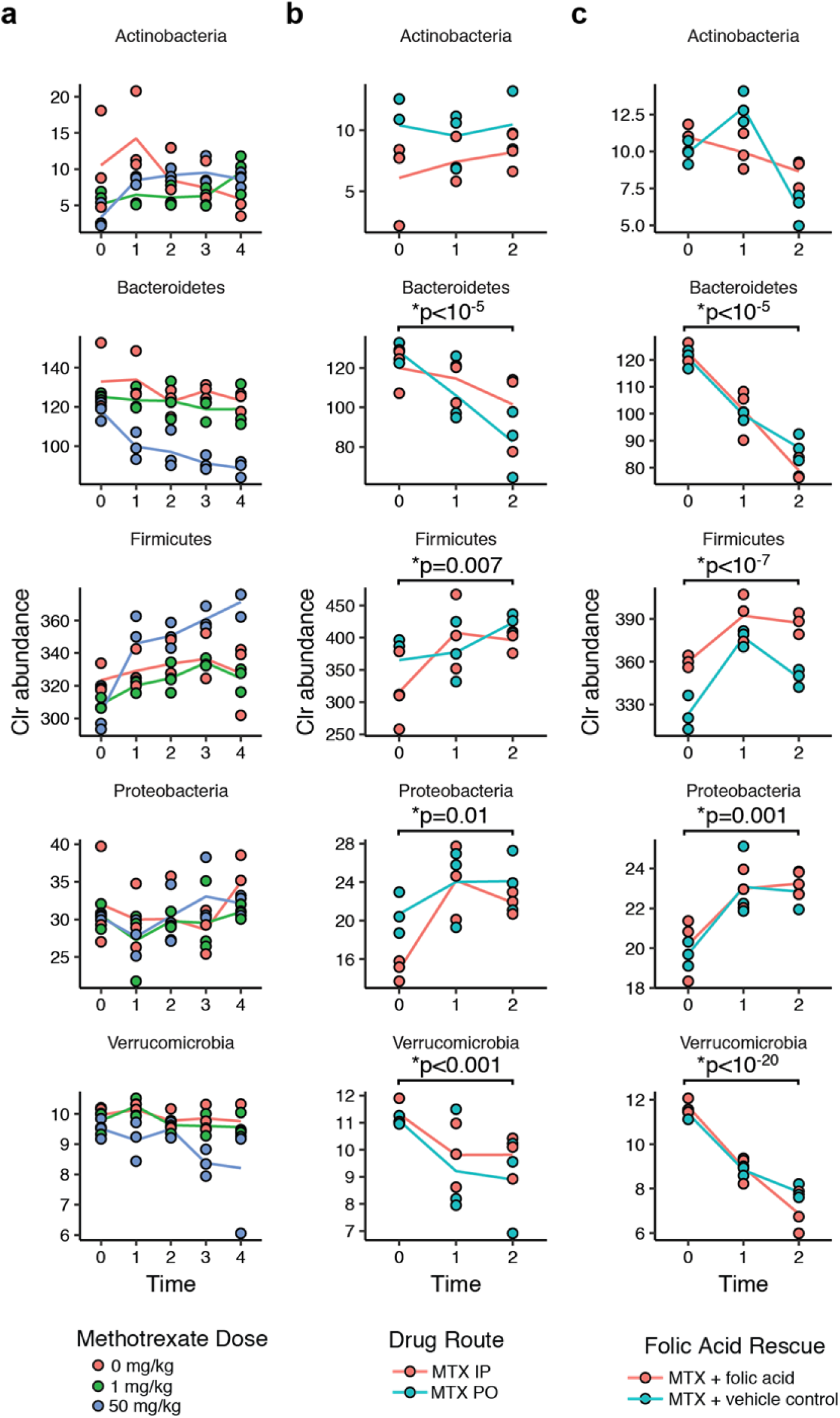
Phylum-level changes are seen in response to MTX treatment. (**a**) Germ-free mice were colonized with a fecal aliquot from a healthy human donor, and treated with either vehicle control, low-dose MTX (1 mg/kg), or high-dose MTX (50 mg/kg) (n=3 per treatment group). Stool samples were collected daily and underwent 16S rRNA gene amplicon sequencing. Shown are the phylum-level trends over time (in days). (**b**) Germ-free mice were colonized with a fecal aliquot from a treatment-naïve rheumatoid arthritis patient and treated with either IP or PO MTX (50 mg/kg) (n=3 per treatment group). Shown are the phylum-level trends. DESeq was used to compare pre- (Day 0) and post-treatment (Day 2) phylum levels among all mice, regardless of treatment group. Significance values are BH-corrected. (**c**) Germ-free mice were colonized with a fecal aliquot from a treatment-naïve rheumatoid arthritis patient and treated with either oral MTX (50 mg/kg) + vehicle control or oral MTX + folic acid (50 mg/kg) (n=3 per treatment group). Shown are the phylum-level trends. DESeq was used to compare pre- (Day 0) and post-treatment (Day 2) phylum levels among all mice, regardless of treatment group. Significance values are BH-corrected. For all plots, colored lines connect the average abundance per treatment group.

**Supplementary Figure 2.**
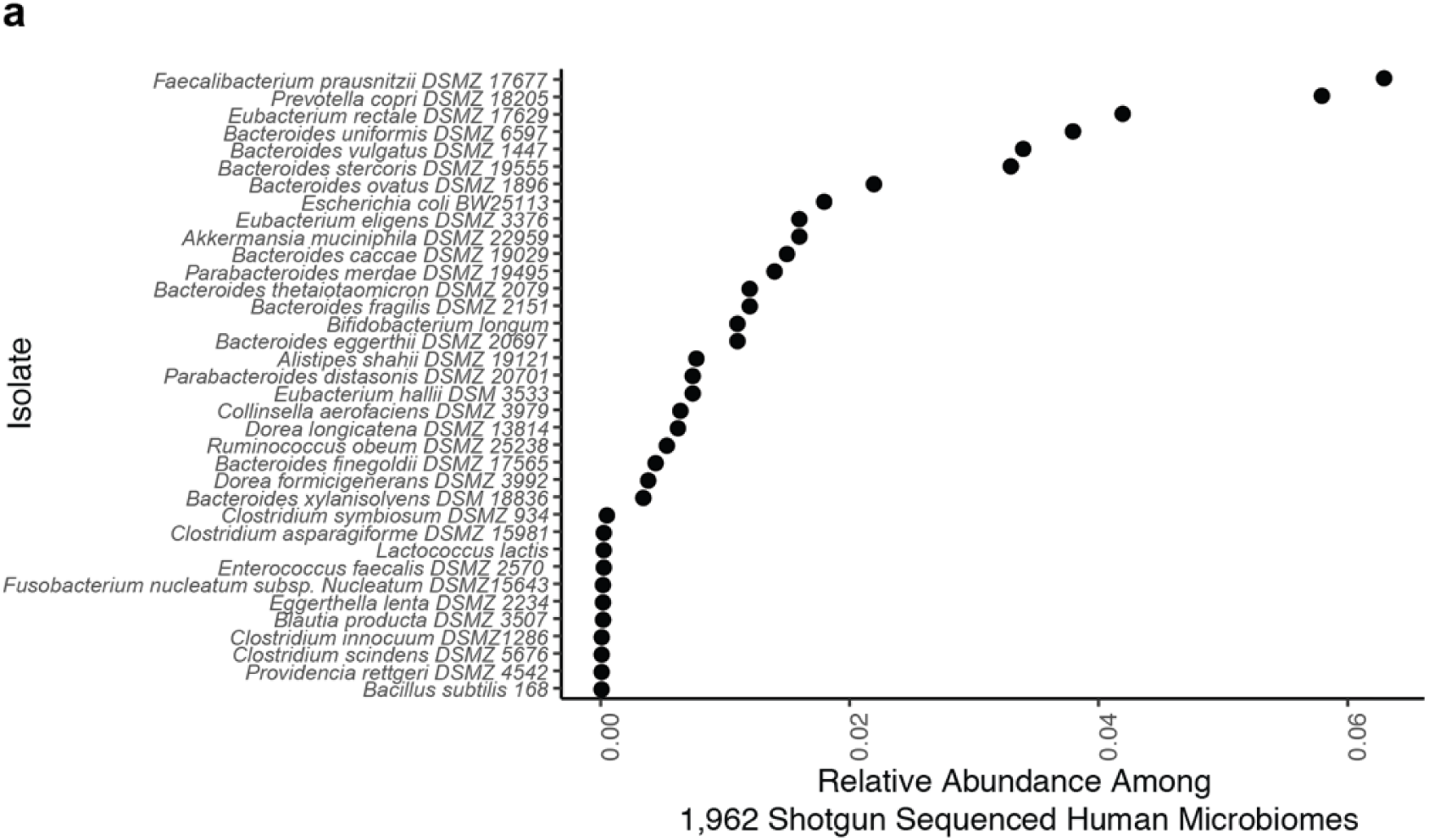
Relative abundance of isolates used in our *in vitro* screen among publicly available gut microbiomes. (**a**) Shown is the mean abundance of each isolate among 1,962 shotgun sequenced microbiomes from healthy and diseased human individuals.

**Supplementary Figure 3.**
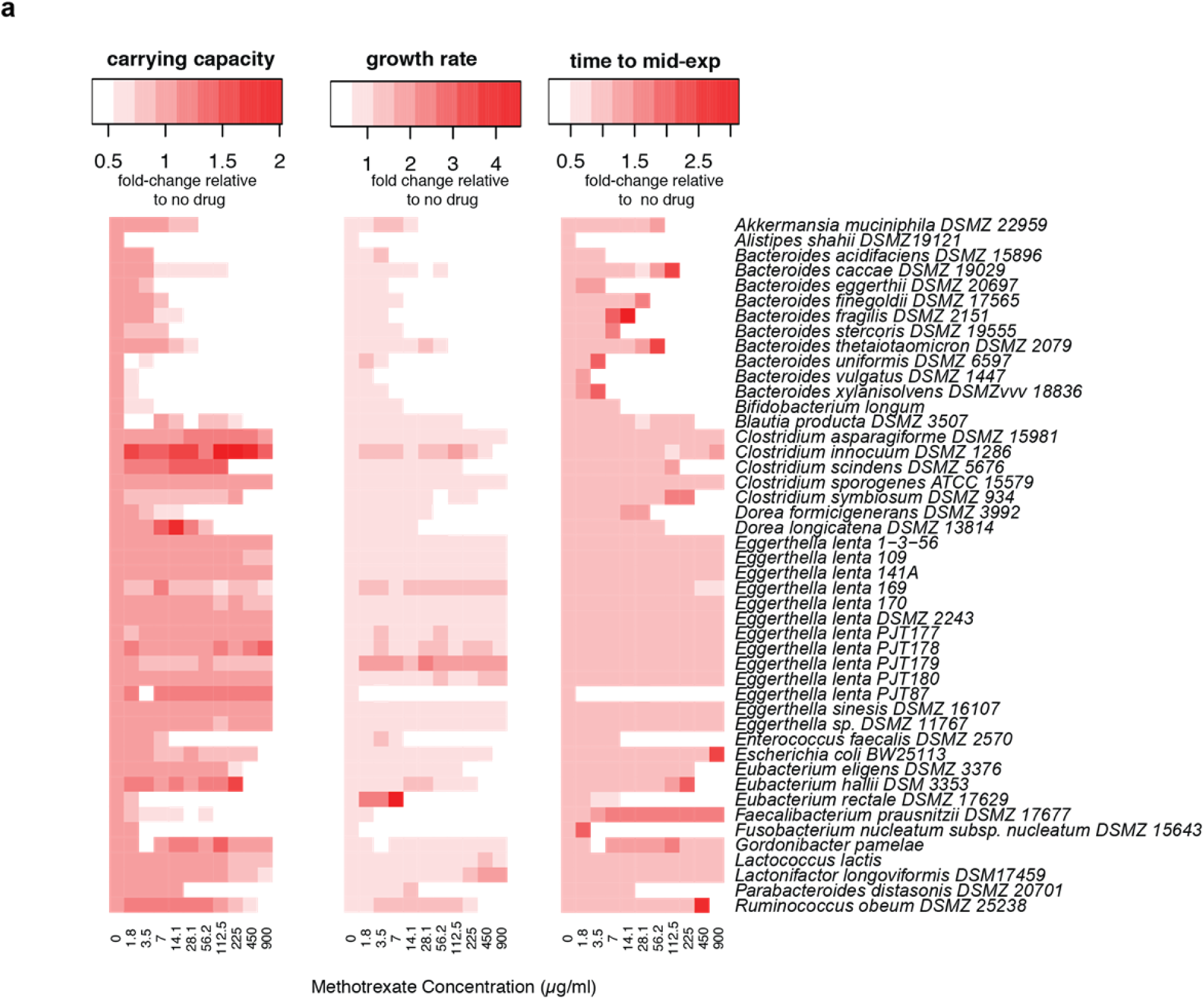
Growth curve parameters are altered by MTX in a dose-dependent manner among human gut bacterial isolates. (**a**) Heatmaps depicting MTX-induced, dose-dependent changes in carrying capacity, growth rate and time to mid-exponential among isolates tested *in vitro*. Changes are relative to parameters measured from growth controls (no drug treatment).

**Supplementary Figure 4.**
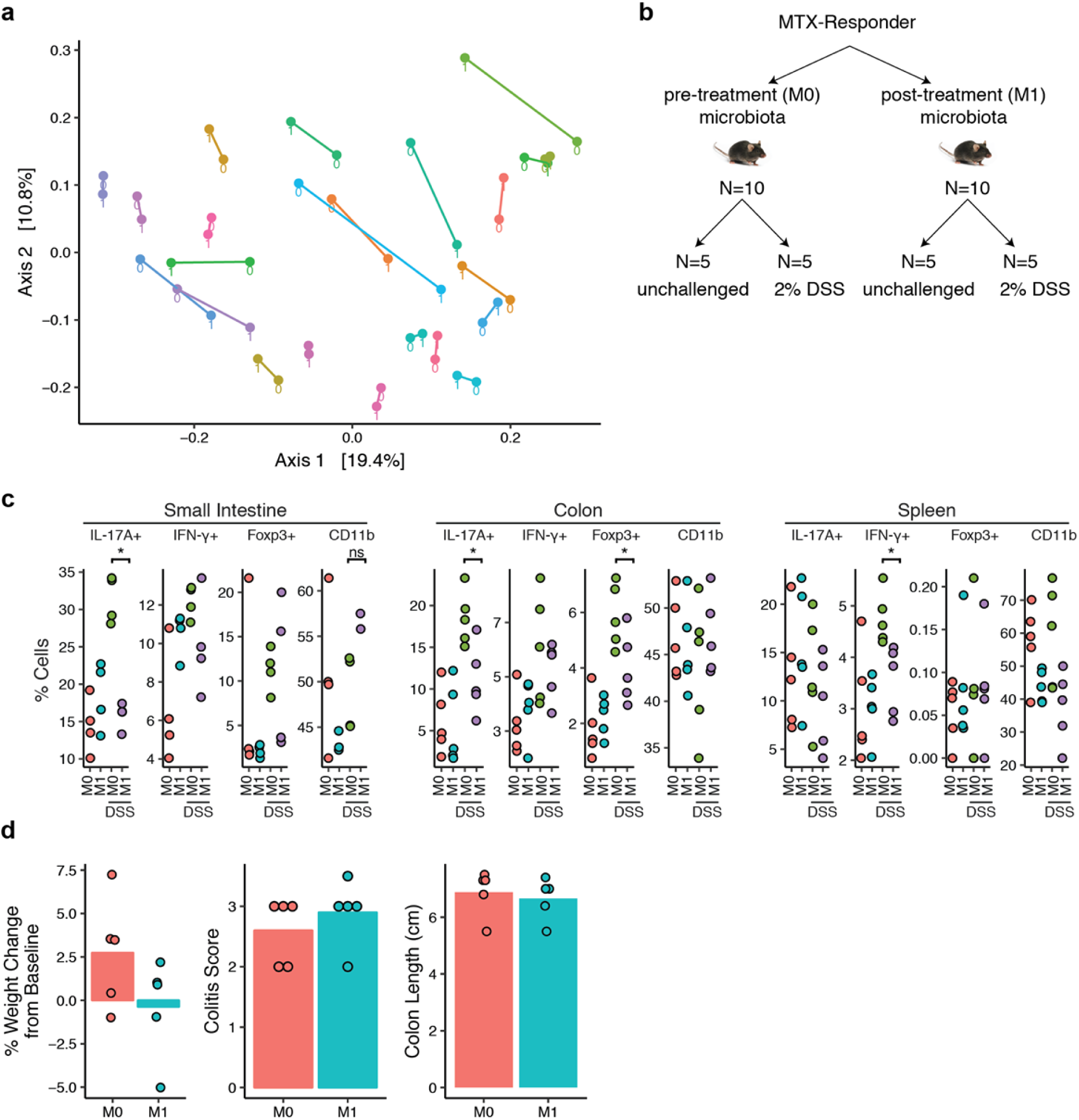
Changes in community composition among rheumatoid arthritis patient microbiomes. (**a**) Principal coordinates analysis (PCoA) of Unifrac distances before (“0”) and 1 month after (“1”) initiation of MTX treatment. Each patient is represented by a unique color and lines connect “0” and “1” time points for that patient. (**b**) Design of a gnotobiotic transplant experiment in which fecal microbiota from a patient before (M0) and 1 month after MTX treatment (M1) were transferred into germ-free C57BL/6J mice. A subset of mice were challenged with dextran sodium sulfate (DSS). (**c**) Percentage of CD11b+ cells or CD3+CD4+ T cells with IL-17A+, IFN-γ +, or FoxP3+ staining from the small intestine and colon lamina propria and spleen of mice transplanted with microbiota from a patient before MTX treatment (“M0”) and 1 month after treatment (“M1”) (N=2-5 per treatment group). Some mice were treated with dextran sodium sulfate (“DSS”) in their drinking water. (**d**) Change in weight, colitis score or colon lengths from mice treated with DSS. Each circle represents a single mouse. Each bar represents the mean value among the mice in each transplant group.

**Supplementary Figure 5.**
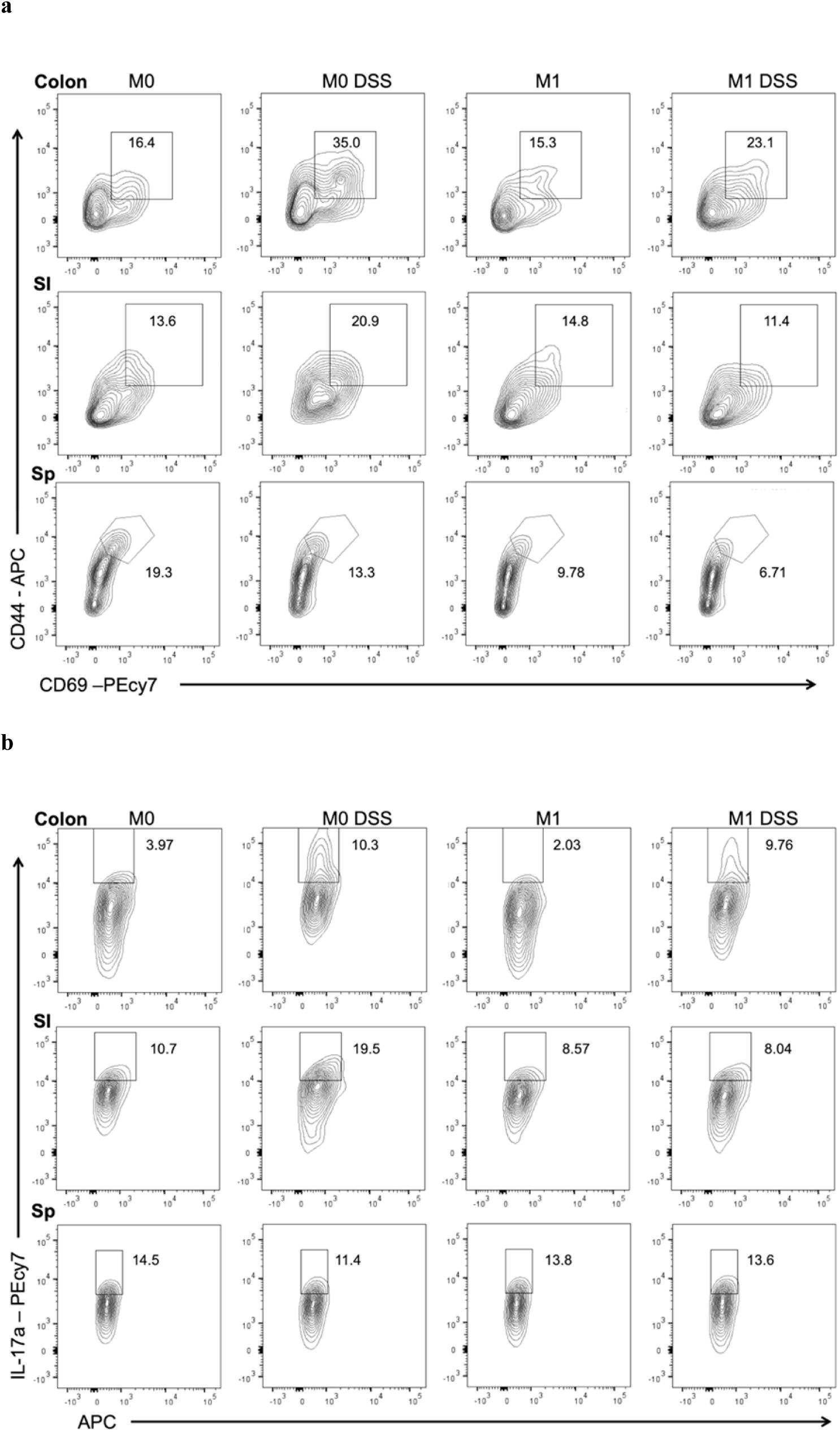
Representative flow plots for immune populations in mice transplanted with microbiota from a patient before MTX treatment (M0) and 1 month after treatment (M1) with or without DSS treatment. (**a**) Colonic and small intestinal lamina (SI) propria lymphocytes as well as splenocytes (Sp) were isolated and stained for flow cytometry analysis. Representative flow plots are shown where the percentage of CD69+ CD44+ cells within the CD3+CD4+ T cells compartment are displayed. (**b**) Representative flow plots of the percentages of Th17 cells as assessed by IL-17A+ within the CD3+ CD4+ compartment.

## Methods

### Gnotobiotic mouse studies

C57BL/6J mice (females, ages 8-16 weeks) were obtained from the UCSF Gnotobiotics core facility (gnotobiotics.ucsf.edu) and co-housed in gnotobiotic isolators for the duration of each experiment (Class Biologically Clean). Mice were colonized with stool from human donors, either a healthy male donor or a treatment-naïve female and male donors with rheumatoid arthritis (as defined by American College of Rheumatology classification criteria^55^). For colonization with a human microbiome, stool was diluted 1:10 g/mL in reduced PBS or saline and homogenized in an anaerobic chamber using pre-equilibrated reagents and supplies. Insoluble material was separated from supernatant by centrifugation at 50*g* for 1 minute. Aliquots of supernatant (200 ul per mouse) were gavaged into mice at least 1-2 weeks before initiation of treatment with MTX. Mice were treated either with saline/PBS, MTX 1 mg/kg or 50 mg/kg daily or folic acid 50 mg/kg. Treatment was carried out either by oral gavage or intra-peritoneal injection. Mice were monitored and weighed daily during treatment. No gross signs of toxicity and minimal-to-no weight loss were observed for the short MTX treatment durations used in this study. Stool samples were collected daily during treatment. Following treatment, mice were euthanized, and contents from the ileum, cecum and colon were collected for sequencing. For dextran sodium sulfate treatment (DSS) (Alfa Aesar, Cat no. 9011-18-1), mice were given 2% DSS (w/v) ad libitum in their drinking water. Mice were monitored for disease progression and weighed daily. Gross signs of toxicity, including hematochezia and weight loss were observed in this study. Stools were scored as follows: 0 = normal stool consistency, 1 = soft stool, 2 = blood in stool, 3 = bloody anus, 4 = prolapsed anus, 5 = moribund/death. All mouse experiments were approved by the University of California San Francisco Institutional Animal Care and Use Committee.

### 16S rRNA gene sequencing of humanized mouse gut microbiota

Aliquots of 108 mouse fecal, ileal, cecal and colon samples (**Supplementary Table 1**) were homogenized with bead beating for 5 min (Mini-Beadbeater-24, BioSpec) using beads of mixed size and material (Lysing Matrix E 2mL Tube, MP Biomedicals) using the digestion solution and lysis buffer of a Wizard SV 96 Genome DNA kit (Promega). The samples were then centrifuged for 10 min at 16,000*g* and the supernatant was transferred to the binding plate. The DNA was then purified according to the manufacturer’s instructions.

16S rRNA gene PCR was carried out as before using GoLay-barcoded 515F/806R primers^56^ according to the methods of the Earth Microbiome Project (earthmicrobiome.org). 2µL of DNA was combined with 25 µL of AmpliTaq Gold 360 Master Mix (Life Technologies) 5 µL of primers (2µM each GoLay-barcoded 515/806R), and 18µL H_2_O. Amplification was as follows: 10 min 95°C, 25x (30s 95°C, 30s 50°C, 30s 72°C), and 7 min 72°C. Amplicons were quantified with PicoGreen (Quant-It dsDNA; Life Technologies) and pooled at equimolar concentrations. Libraries were then quantified (NEBNext Library Quantification Kit; New England Biolabs) and sequenced with a 600 cycle MiSeq Reagent Kit (251×151; Illumina) with ∼10% PhiX.

### 16S rRNA amplicon analysis of mouse and human fecal samples

Reads were demultiplexed using QIIME^57^ v1.9.1 (split_libraries_fastq.py) before denoising and processing with DADA2^58^ v1.1.5 under MRO v3.2.5. Taxonomy was assigned using the DADA2 implementation of the RDP classifier^59^ using the DADA2-formatted RDP v14 training set (benjjneb.github.io/dada2/assign.html). A phylogenetic tree was constructed using DECIPHER v2.8.1 and PHANGHORN v2.4.0 as described in Callahan et al^60^. Diversity metrics were generated using Vegan v2.4-3 and Phyloseq v1.20.0^61^ with principal coordinate analysis (PCoA) carried out with Ape v4.1. Analyses were carried out on either: (1) centered log2-ratio (clr) normalized taxonomic abundances calculated as A_clr_=[log_2_(A_1_/g_a_), log_2_(A_2_/g_a_),… log_2_(A_n_/g_a_),], where A is a vector of non-zero read counts and g_a_ is the geometric mean of all values of A, or (2) relative abundance calculated as proportion of reads. ANOSIM and PERMANOVA were used to detect changes in community composition using counts from rarefied data and Bray-Curtis distances. DESeq2^62^ was used to determine differentially abundant taxa on raw count data. Significance testing of longitudinal trends was determined using generalized mixed effects models using the cplm package^22,63^ (v. 0.7-7) on clr normalized values. For each sample, fastq files are available in NCBI’s Sequence Read Archive (SRA), accession number SRP5125967.

### Quantitative PCR for 16S copy number determination

Quantitative PCR (qPCR) of total 16S rRNA gene copies was carried out in triplicate 10µL reactions with 200nM 891F(5’-TGGAGCATGTGGTTTAATTCGA-3’)/1003R(5’-TGCGGGACTTAACCCAACA-3’) primers using a BioRad CFX384 thermocycler with iTaq^TM^ Universal Probes Supermix (BioRad 1725132) and probe 1002P ([Cy5]CACGAGCTGACGACARCCATGCA[BHQ3]) according to the manufacturer’s instructions and an annealing temperature of 60°C. Absolute quantifications were determined based against a standard curve of 8F/1542R amplified from purified bacterial DNA. Reactions were performed in triplicate and mean values were taken for further downstream analyses. Absolute bacterial abundance was derived by adjustments for dilutions during DNA extraction, normalization, and PCR reaction preparation dividing by the total fecal mass used for DNA extraction in grams.

### *In vitro* bacterial growth studies

The isolates used in this study are shown in **Supplementary Table 9**. Each of these strains was obtained from the Deutsche Sammlung von Mikroorganismen und Zellkulturen (DSMZ) culture collection. A single colony of each isolate was subcultured in Bacto Brain Heart Infusion (BD Biosciences, 37 g/L) supplemented with *L*-cysteine-HCl (0.05%, w/v), menadione (1 µg/mL), and hemin (5 µg/mL) (referred to hereafter as BHI+) for 48 hours in an anaerobic chamber (Coy Laboratory Products) at 37°C with an atmosphere composed of 2-3% H_2_, 20% CO_2_, and the balance N_2_. This subculture diluted down to an OD600 of 0.08-0.1, which was then further diluted 100-fold, and then used to inoculate a microtiter plate with 2-fold serial dilutions of MTX concentrations ranging from 0 – 900 μg/ml. Plates were incubated at 37°C with shaking in an Eon Microplate Spectrophotometer (BioTek Instruments, Inc) over a 48 to 72-hour period in the anaerobic chamber. Growth was monitored every 15 minutes at OD600 and corrected for background (no growth control). Data were exported using the Gen5 (v 2.0) software. The minimal inhibitory concentration (MIC_90_) was measured as the lowest concentration of MTX that resulting in >90% growth inhibition after 48 hours of incubation. Growth parameters (carrying capacity, time to mid-exponential and growth rate) were calculated in R using GrowthCurveR^64^ (v 0.2.1). Determination of dose-dependency was done by fitting a linear regression with methotrexate concentration as the independent variable and the estimated growth parameter the dependent variable using lm from the stats package (v 3.5.1). Heatmaps of these parameters were generated using gplots (v 3.0.1), with each parameter normalized to the parameter derived from growth control curves (i.e. no drug present).

### Tree Construction

Ribosomal sequences for each isolate were extracted from the Greengenes^65^ database (May, 2013). Sequences were imported into UGENE^66^ (v 1.31.0), and aligned using MUSCLE^67^. Gaps occurring in > 50% of sequences were removed, and a maximum likelihood tree was generated using PhyML^68^ with 100 bootstraps and the GTR substitution model.

### Abundance Quantification of Bacterial Isolates from Shotgun Sequencing Data

Bacterial abundances were quantified using data from Metaquery^30^, a web-based application that provides taxanomic abundances from >1,900 publicly available human gut metagenomes. For each isolate, we queried the “metaphlan2” database and recorded the mean abundance value.

### Predicted concentration of MTX in the GI tract

The predicted concentration of MTX in the proximal GI tract was estimated by taking the oral dose used for rheumatoid arthritis (25 mg) and dividing it by 250 ml^31^, giving a concentration of 100 μg/ml or 220 uM.

### MTX Treatment for RNA-Seq

The bacterial strains used in RNA-Seq are given in **Supplementary Table 9.** Genomes are available in NCBI’s GenBank Assembly database (see **Supplementary Table 9** for accession numbers). Culture media was composed of BHI+ and allowed to equilibrate in an anaerobic environment prior to use. Briefly, bacteria were cultured in BHI+ at 37 °C in an anaerobic chamber. Cultures for each isolate were grown to mid-exponential (achieving an OD600 ∼ 0.5), aliquoted into triplicates, treated for 30 minutes with either DMSO or MTX 100 μg/ml, and then removed from the anaerobic chamber. For *C. asparagiforme*, cultures were incubated for 4 and 20 hours as well. Cultures were centrifuged at 2000 rpm for 10 min at 4 °C to facilitate removal of supernatant, and the remaining bacterial pellet was flash-frozen in liquid nitrogen.

### Total RNA extraction

Each bacterial pellet was incubated with 1 ml of Tri reagent (Sigma Aldrich, catalog #: T9424) at room temperature for 10 minutes. The cell suspension was transferred into Lysing Matrix E tubes (MP Biomedicals, 116914050), and homogenized in a bead-beater (Mini-Beadbeater-24, BioSpec) for 5 minutes at room temperature. The sample was incubated with 200 uL of chloroform at room temperature for 10 minutes, followed by centrifugation at 16,000 × *g* for 15 minutes at 4 °C. Next, 500 uL of the upper aqueous phase was transferred into a new tube and 500 uL of 100% ethanol was added. To isolate RNA, we used the PureLink RNA Mini Kit (Life Technologies, catalog #: 12183025). This mixture was transferred onto a PureLink spin column and spun at ≥ 12,000 × *g* for 30 seconds. The column was washed with 350 ul of wash buffer I as described in the PureLink manual. The column was incubated with 80 ul of PureLink DNase (Life Technologies, catalog #: 12185010) at room temperature for 15 minutes, and washed with 350 ul of wash buffer I. The column was washed with wash buffer II twice as described in the PureLink manual. Total RNA was recovered in 50 ul of RNAase-free water. A second round of DNAse treatment was undertaken. The RNA was incubated with 6 ul of TURBO DNAse (Ambion, ThermoFisher, catalog #: AM2238) at 37°C for 30 minutes. To stop the reaction, 56 ul of lysis buffer from the PureLink kit and 56 ul of 100% ethanol was added to the sample and vortexed. This suspension was transferred onto a PureLink column, and washed once with 350 ul of wash buffer I and twice with 500 ul of wash buffer II. The RNA was recovered in 30 ul of RNAse-free water.

### rRNA depletion, library generation, and RNA sequencing

Total RNA was subjected to rRNA depletion using Ribo-Zero Bacterial rRNA Depletion (Illumina, catalog #: MRZB12424), following the manufacturer’s protocol. RNA fragmentation, cDNA synthesis, and library preparation proceeded using NEBNext Ultra RNA Library Prep Kit for Illumina (New England BioLabs, catalog #: E7530) and NEBNext Multiplex Oligos for Illumina, Dual Index Primers (New England BioLabs, catalog #: E7600), following the manufacturer’s protocol. All samples were single-end sequenced (1×50 bp) using an Illumina HiSeq2500 platform (High Output, v4 chemistry) at UCSF’s Institute for Human Genomics. For each sample, fastq files are available in NCBI’s Sequence Read Archive (SRA), accession number SRP5125967.

### RNA-Seq analysis

Reads were mapped to reference genomes using Bowtie2^69^. HTSeq (v 0.8.0) was used to count the number of transcripts mapping to genes^70^. Differential gene expression was assessed using DESeq2^62^. KEGG Pathway enrichment was carried out using clusterProfiler^71^ (v3.4.1).

### Human RA patient samples acquisition

Consecutive patients from the New York University Langone Medical Center’s rheumatology clinics and offices were screened for the presence of RA based on ACR criteria^55^. After informed consent was signed, each patient’s medical history (according to chart review and interview/questionnaire), diet, and medications were determined. A screening musculoskeletal examination and laboratory assessments were also performed or reviewed. All RA patients who met the study criteria were offered enrollment. The criteria for inclusion in the study required that patients meet the American College of Rheumatology/European League Against Rheumatism 2010 classification criteria for RA^55^, including seropositivity for rheumatoid factor (RF) and/or anti–citrullinated protein antibodies (ACPAs), and that all subjects be age 18 years or older. New-onset RA was defined as disease duration of a minimum of 6 weeks and up to 6 months since diagnosis, and absence of any treatment with disease-modifying anti-rheumatic drugs (DMARDs), biologic therapy or steroids (ever). The exclusion criteria applied to all groups were as follows: recent (<3 months prior) use of any antibiotic therapy, current extreme diet (e.g., parenteral nutrition or macrobiotic diet), known inflammatory bowel disease, known history of malignancy, current consumption of probiotics, any gastrointestinal tract surgery leaving permanent residua (e.g., gastrectomy, bariatric surgery, colectomy), or significant liver, renal, or peptic ulcer disease. This study was approved by the Institutional Review Board of New York University School of Medicine protocols #09-0658 and # as previously published^43^. All new onset rheumatoid arthritis (NORA) patients (n=30 patients, 60 stool samples) were recruited using established protocols from a previously described study^43^. Patients received oral MTX at standard of care doses as prescribed by their treating rheumatologists. Stool samples were collected at baseline and 1 month after MTX initiation and metadata were obtained at baseline and 4 months after therapy initiation. Clinical and demographic data was de-identified and recorded in RedCap by the designated study personnel. Clinical responder status (MTX-R) was defined *a priori* as any NORA patient whose DAS28 score was greater than 2 at baseline and improved by at least 1.8 by month 4 post-treatment. DNA was extracted from human fecal samples (n=23 patients, 46 stool samples) as we have previously described^43^, using the MoBio Powersoil DNA extraction kit, based on cell membrane disruption by high-speed shaking in the presence of beads. The V4 hypervariable region of bacterial 16S ribosomal RNA (rRNA) was performed using a MiSeq Illumina platform (150□ bp read length, paired□ end protocol) at the New York University Genome Technology Center as previously described^72^. For each sample, fastq files are available in NCBI’s Sequence Read Archive (SRA), accession number SRP5125967.

### *Ex vivo* incubation of RA patient stool samples

All work was carried out in an anaerobic chamber. For each patient, stool was aliquoted into a pre-equilibrated cryovial, diluted in reduced PBS at 10 ml per 1 gram of stool, and vortexed to homogenize the sample. The sample was spun at ∼20*g* for 1 minute on a mini-centrifuge to facilitate settling of sediment, and the sediment-free supernatant was then aliquoted into a new pre-equilibrated cryovial for evaluation of *ex vivo* growth. Growth was evaluated by inoculating liquid BHI with 1:50 dilution of this fecal slurry, with OD600 readings performed every 15 minutes for 48 hours with a 2-minute shake prior to each reading. Samples were treated with MTX 100 μg/ml or an equal volume of DMSO at time zero. Each patient’s fecal slurry and treatment was evaluated in quadruplicate. Growth curves were averaged by treatment and individual, and growth parameters were estimated using the GrowthCurveR package (v 0.2.1). Paired Student’s *t*-tests were used to determine changes in growth parameters.

### Lamina Propria Lymphocyte Isolation

Lamina propria lymphocytes (LPLs) were isolated with slight modifications of previously described methods^73-75^. In brief, small intestinal (SI) Peyer’s patches were removed and colons and SI tissue were splayed longitudinally with mucus removed by scraping and stored in complete RPMI (10% fetal bovine serum, 100 units per ml penicillin and streptomycin, β-mercaptoethanol, glutamate, sodium pyruvate, HEPES and non-essential amino acids). Supernatants were removed by filtering through a 100μM filter, and remaining tissue incubated in 1X HBSS (without Ca^2+^ and Mg^2+^) containing 5 mM EDTA (Promega) and 1 mM DL-Dithiothreitol (DTT) (Bioplus chemicals) for 45 min at 37°C on a shaker. Supernatant was removed by filtering through a 100 μM filter, and remaining tissue was incubated for 45 min (colon) or 35 min (SI) at 37°C on a shaker in a solution containing 1X HBSS containing 5% (v/v) fetal bovine serum (GIBCO heat inactivated), 1 U/ml Dispase (Sigma), 0.5 mg/ml Collagenase VIII (Sigma), and 20 μg/ml DNaseI (Sigma). The supernatant was filtered over a 40 mm cell strainer into ice-cold sterile 1X PBS. Cells were subjected to a Percoll (VWR) gradient (40%/80% [v/v] gradient) and spun at 2000RPM for 20 min with no brake and no acceleration. Cells at the interface were collected, washed in PBS and prepared for flow cytometry analysis.

### Flow Cytometry

Lymphocytes were isolated from the colonic and small intestinal lamina propria as described above. Spleen cells were prepped through gentle mashing with a syringe plunger. Spleen cells were treated with 1X RBC Lysis Buffer (Biolegend) to lyse and remove red blood cells. Surface staining for lymphocytes was done in staining buffer (1X HBSS (Corning) supplemented with 10 mM HEPES (Cellgro), 2 mM EDTA (Promega), and 0.5% (v/v) fetal bovine serum (GIBCO heat inactivated)) for 20 min at 4°C. Cells were then washed twice in supplemented 1X HBSS and enumerated via flow cytometry. The following antibodies were used: anti-CD3 (17A2, Invitrogen), anti-CD4 (GK1.5, Biolegend), anti-CD69 (H1.2F3, Biolegend), anti-CD11b (M1/70, Biolegend), and anti-CD44 (IM7, Tonbo biosciences). For intracellular staining, cells were first stimulated with ionomycin (1000 ng/ml), PMA (50 ng/ml), and Golgi Plug (1 μl/sample) (BD Bioscience) overnight at 37°C. Stimulated cells were stained with LIVE/DEAD Fixable Aqua Dead Cell Stain Kit (Thermo Fisher) CD3+ CD4+ cells were assessed within the live population. Cells were surface stained, washed, and then fixed/permeabilized in 100μl Perm/Fix buffer (BD Bioscience). Cells were washed twice in Perm/Wash buffer (BD Bioscience) and then stained for intracellular cytokines with the following antibodies: anti-IFN-γ (XMG1.2, Millipore), anti-IL-17A (ebio17B7, Invitrogen), anti-Foxp3 (150D, Biolegend). Cells were washed twice in Perm/Wash buffer and then placed in staining buffer for flow cytometry analysis. Gating cell populations was done using isotype and single stain controls. Representative gating strategies are provided in **Supplementary Fig. 5**. These data were collected with a BD LSR Fortessa and analyzed with FlowJo software.

### Data Availability

All sequencing data generated in the preparation of this manuscript has been deposited in NCBI’s Sequence Read Archive (SRA), accession number SRP5125967

## Code Availability

Code used in the analyses presented here are publicly available as described in Callahan et al^60^ and upon request.

